# Systematic Regional Bias is Widespread in ChIP-seq

**DOI:** 10.64898/2026.05.10.724164

**Authors:** Oliver Hughes, Gabriel Foley, Brad Balderson, Michael Piper, Mikael Bodén

## Abstract

Robust and reproducible results are essential for confident scientific analysis. We demonstrate that transcription factor (TF) Chromatin Immunoprecipitation coupled with sequencing (ChIP-seq) suffers from systematic bias that may threaten its reproducibility: 80% of 200+ condition-matched, dual-replicate experiments in ENCODE contain genomic regions of systematic bias. We observe this regional bias even between replicates produced within the same experiment, resulting in thousands of unreplicated peaks, which often contain valuable biological data. We provide evidence that regional bias may lead to qualitative differences in TF biology inferred by different experiments; we discovered eight TFs with binding activity in compact chromatin that was identified by one experiment, yet systematically absent from others. To mitigate the effects of bias, we derive simple but effective metrics to quantify the quality of data within biased regions and demonstrate that they can be used for the robust integration of data from multiple experiments.

## INTRODUCTION

Chromatin immunoprecipitation paired with next generation sequencing (ChIP-seq) is a technology central to the study of transcription factors (TF). Designed to detect TF binding *in vivo*, ChIP-seq has been used to build genome wide maps of their DNA binding sites within a diverse array of cells, tissues and organisms^1^. The scientific community has developed a wide array of computational tools to assess the quality of ChIP-seq data, and large consortia such as ENCODE^2^ provide a central repository for thousands of experiments produced according to uniform guidelines.

Tools such as IDR^3^, MSPC^4^ and others^5,6^ employ “consensus” filtering, which assumes that “peaks” (loci that are statistically enriched for ChIP-seq reads) are identified consistently across a set of replicates or experiments, are of higher quality than those discovered less frequently. Whilst intuitive, that consensus peaks are of inherently higher quality has not been thoroughly confirmed in the ChIP-seq literature.

Indeed, during the development of the tool ChIP-R^7^, we discovered that inconsistent peaks often represent genuine binding events, and restricting peaks to high levels of consensus sometimes results in poorer, not higher quality data. Upon investigating this phenomenon, we made a startling discovery: condition-matched ChIP-seq experiments feature regions of severe bias, occupying megabase scale domains and influencing the reproducibility of tens of thousands of peaks.

Previous work in ChIP-seq has catalogued large contiguous regions prone to generating spurious signal across multiple experiments, with the ENCODE Blacklist Regions^8^ being the most well-known curation. Despite this, little is known about systematic variation *between* ChIP-seq experiments, or even replicates.

Thus, the discovery of systematic regional bias exposes a longstanding gap in the literature. In technologies such as RNA-seq, systematic biases or “batch effects” are well recognised, with a vast array of tools available to mitigate their tendency to drive spurious results and obscure real biological relationships^9,10^.

With no equivalent resources to detect or address bias in ChIP-seq, the extent to which regional bias impacts the ability of researchers to make inferences about TF biology remains unknown.

We addressed this gap by developing the first methodology capable of detecting systematic biases between ChIP-seq replicates or matching experiments: Hidden Markov Models (HMMs)^11^ are used to detect shifting patterns of agreement across replicates (termed “overlap patterns”) and cluster sequences of neighbouring peaks into a set of regional patterns (referred to as “states”). We detected regional bias in 80% of condition-matched human TF ChIP-seq experiments and 50% of within-experiments replicates, which confirmed that regional bias is a pervasive issue in ChIP-seq.

In this paper, we answer three key questions: I) Do regions of bias (or simply low consensus regions) contain biologically meaningful information? II) What metrics can distinguish high from low quality regions in the absence of consensus filtering? III) Can systematic regional bias lead to fundamental differences in the biology inferred for the same TF across different experiments?

We used TF-specific motif analysis to answer I), establishing that peaks within regions of bias often represent genuine TF activity. Strikingly, regional consensus is inconsistently associated with data quality overall, suggesting that consensus filtering - employed by tools like ChIP-R and IDR - may not be an appropriate strategy for integrating ChIP-seq replicates or experiments. To address II), we demonstrate that low quality regions of bias between experiments are characterised by systematically weak ChIP-signal relative to high quality regions, allowing for researchers to identify regions of peaks representing valid binding sites, even if they are systematically under-discovered in some experiments. Despite this success, ChIP-signal was a poor predictor of data quality for regions of bias between replicates (within experiments), and thus such regions must still be treated with caution by researchers.

Finally, we answered III): By accounting for bias between experiments, we revealed eight unique TFs that exhibit binding in closed chromatin. We uncovered robust evidence linking bias in less accessible chromatin to fundamental differences between experiments in the ability to extract reads from closed chromatin, suggesting that technical variability can have a profound effect on the information contained between different ChIP-seq experiments. Peaks in regions of less accessible chromatin featured robust TF-specific motif enrichment and featured amongst the strongest ChIP-signal in every dataset, yet only one TF (FOXA2) is broadly recognised as playing a functional role in closed chromatin^12^.

We suggest that regional bias, resulting from technical variability, may obscure fundamental aspects of the biology of otherwise well studied TFs. Given the high frequency of regional bias in ChIP-seq data, careful integration of data from multiple experiments may be necessary to obtain comprehensive profiles of TF biology.

## RESULTS

### A Model of Systematic Regional Bias

We detect regional bias via statistical changes in the sequence of overlap patterns between a set of replicates (defined in terms of a “peak union”, see Methods and Materials). For example, a recurring sequence of the overlap pattern “T T F F” represents a region where peaks are systematically absent (F for False, as opposed to T for True) from the last two (of four) replicates. We employ HMMs to perform unsupervised discovery of such patterns by clustering segments of the sequence of overlaps into a set of discrete states. The advantage of this approach is that it obviates the need to define regional boundaries *a-priori*, instead, the HMM enables purely data driven discovery of regional patterns operating at any spatial scale. A graphical overview of this process can be seen in Figure 1, along with a demonstration of regional bias taken from real data.

**Figure 1:**
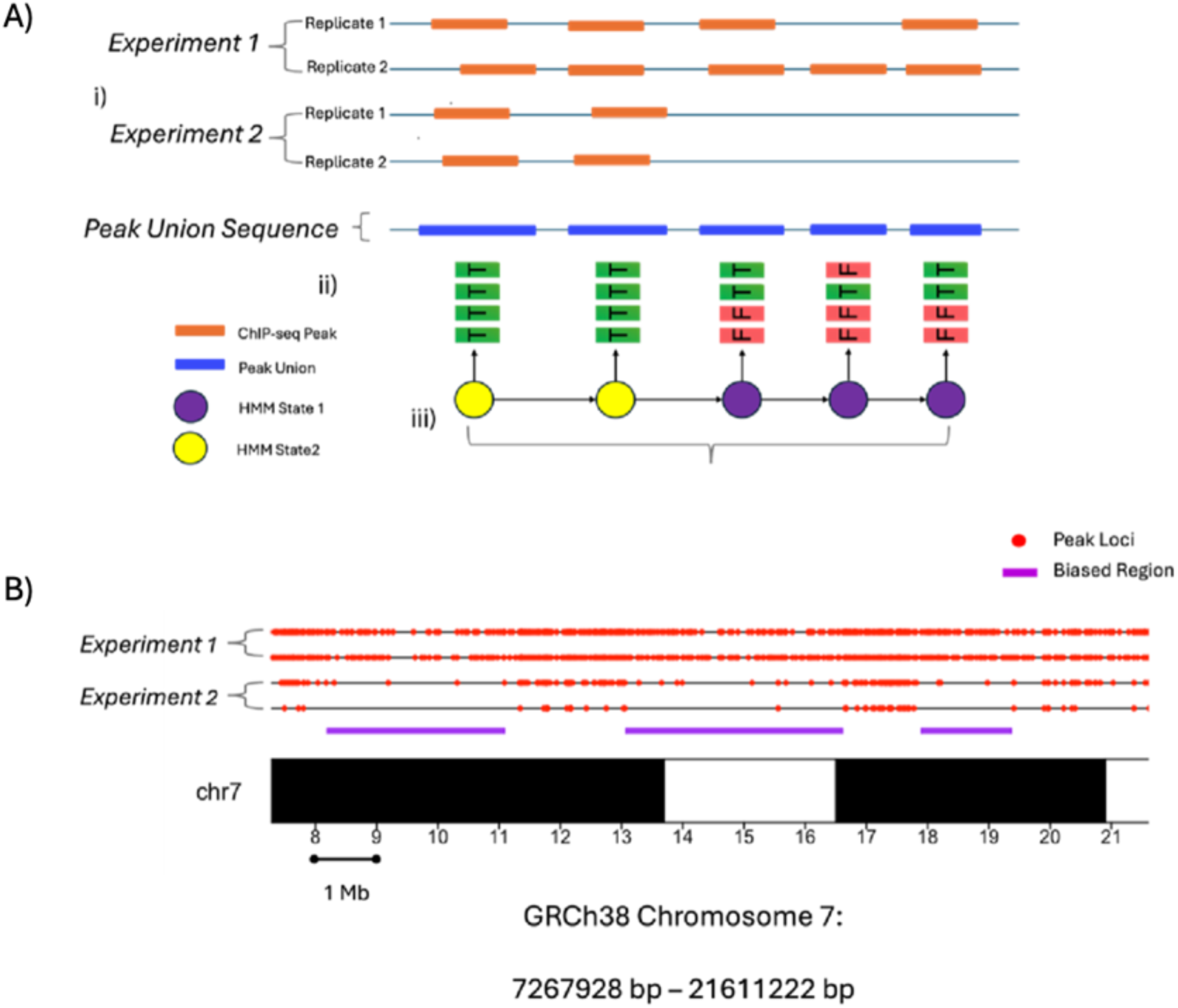
A) Graphical overview of modelling regional bias. i) Peaks from all replicates are merged into a sequence of non-overlapping unified intervals (peak unions); ii) The combination of replicates contributing to a peak union are encoded as T/F tuples; iii) The HMM models change in the distribution of overlap patterns as changes in a sequence of latent states. B) Three biased regions on chromosome 7 of experiment set CEBPB-A549, state annotations of adjacent unbiased regions not shown.

### Regional Bias Detected Between 80% of Matched ENCODE Experiments, Half of Replicates from the Same Experiment

We quantified the prevalence of regional bias for all pairs of matching human TF ChIP-seq experiments (experiment sets) in ENCODE (n=229), and to the replicates within the remaining unmatched human ChIP-seq experiments (replicate sets) (n=1792). We applied the HMM to these datasets, varying the number of latent states between 1-6 to identify the optimal number of biased patterns in each dataset. State selection using the BIC^13^ revealed that 184/229 (80%) of experiment sets featured an optimal number of states >1, signalling the presence of regional bias. The unmatched experiments revealed that even replicates within experiments are subject to systematic regional bias, with 49% (884/1792) of experiments optimally having more than one state when their replicates were compared.

To validate our approach, we ran HMMs on randomised datasets to test if the BIC would erroneously identify systematic bias in random sequences. We constructed random datasets by permuting the order of overlap patterns in each dataset, which ensures that the random datasets were identical in size and composition to the real data.

Across 2021 random datasets (experiment and replicate sets combined), the BIC never returned an optimal number of states >1, showing that our approach has a negligible false positive rate. This result provides robust evidence that the latent states discovered by the HMMs represent genuine systematic bias between experiments.

The biased patterns between experiment sets often represented regions where only one experiment identified the majority of peaks; 45% of experiment sets featured a latent state where >70% of peaks were identified in one experiment alone, representing recurring regions dominated by a single experiment.

A large proportion of our ENCODE experiment sets (52%, 119/229) featured multiple matching experiments performed by the same laboratory. We tested if within-laboratory experiment sets were less likely to feature regional bias than between-laboratory sets, and found no significant difference in the frequency of regional bias (𝜒^2^ =0, *p* = 1) when experiments were performed by the same laboratory 81% (74/91) than when they were produced by different laboratories 80% (111/138).

### Biased Regions Occupy Vast Regional Domains, Influencing the Reproducibility of Thousands of Peaks

We found that segments of peaks annotated to experiment biased states spanned large, megabase scale regions, reoccurring multiple times throughout the genome. The most biased state in each experiment set occupied a median of 52 discrete genomic regions (Interquartile range (IQR) : 98, 27-125), each spanning a median of 8.04 Mb (IQR: 29.63, 1.48-31.11) and containing a median of 8670 peaks (IQR: 13081, 2658.5-15739.5) of which a median of 76% (IQR: 33%, 59%-86%) were exclusive to one experiment.

We also discovered that biased regions were non randomly distributed throughout the genome, with the odds of an overlapping state being the most biased in its experiment set differing between Giemsa staining categories^14^ (*F*=15.94, p < 1e-04, permutation test).

Dark stained, heterochromatin rich cytoband regions of the genome contained the highest enrichment for biased regions (see Figure 2). On average, regions overlapping the darkest staining category (gpos100) had 15% greater odds of being the most biased state in its respective experiment than light coloured, euchromatin bands (gneg) (P-val). Additionally, 22% (18/81) of gpos100 stained bands were amongst the top 10% most bias-enriched cytobands compared to only 8% (34/413) of gneg stained bands. This means that there is a 2.75 fold enrichment for dark stained cytobands amongst the top 10% bias hotspots across the genome.

**Figure 2:**
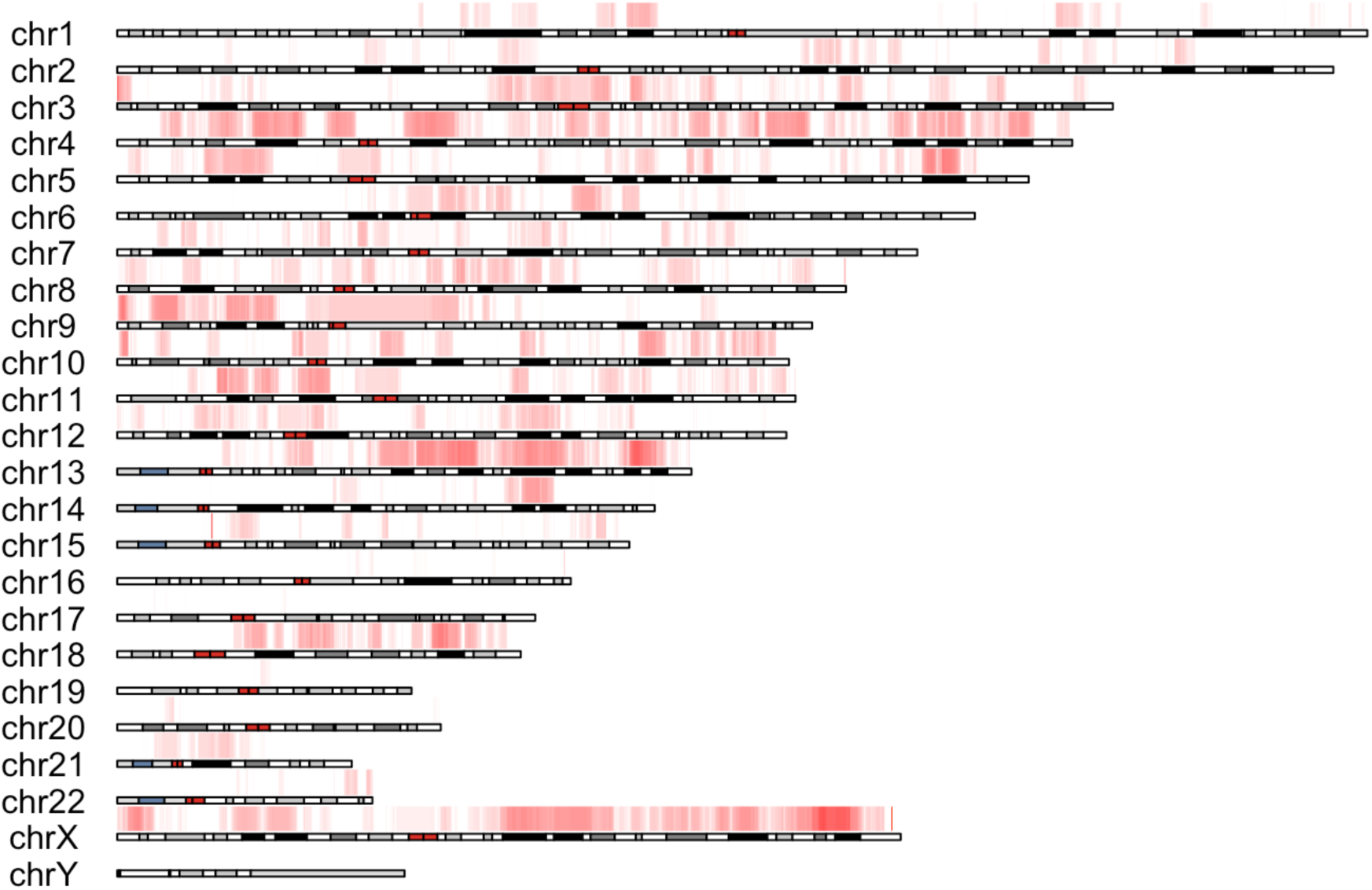
Regions of Bias Between Experiments are statistically enriched in darker stained cytobands: A heatmap displaying the statistical enrichment of regions of bias between ENCODE experiments across the human genome. For each experiment set, the maximum “Experiment biased” regions were defined for each experiment set as the HMM state with the greatest number of peak unions identified exclusively by a single experiment. The human genome assembly GRCh38 was divided into 100Kb bins, and for each bin, the proportion of overlapping states that were the maximum biased states in their respective experiment sets was calculated. Plotted values for each bin are proportional to the odds that that an overlapping state is the most biased state in its experiment set, P(Maximum Biased State)/(1-P(Maximum Biased State).

### Regional Bias is Frequent Across Commonly used Peak Callers

Regional bias is highly frequent across ENCODE, however, ENCODE provides data for peaks called by only one peak caller, MACS2^15^. We replicated our analysis using different peak callers to assess whether regional bias is consistent across different computational tools.

We randomly sampled 90 experiment sets from ENCODE and re-called peaks using the peak callers MACS2, GEM^16^ and SISSRs^17^, each chosen to maximise the diversity of computational approaches across peak callers.

The frequency of regional bias was similarly high across peak callers; regional bias was detected by the HMM in 80/90 (89%), 61/90 (68%) and 73/90 (81%) experiment sets for MACS2, GEM and SISSRs respectively. Of the 90 experiments, 32 contained regional bias across all three peak callers when restricted to a common set of peaks (See Methods and Materials).

To quantify the similarity of patterns of bias between peak callers, we first defined a common set of peak loci by taking the intersection between the peak union sequences generated by peaks called with each peak caller. This results in a set of peak loci unanimously discovered by each peak caller, yet overlap patterns between replicates may differ between each peak caller. We aligned states across HMMs (from each peak caller) by finding overlapping Viterbi annotations and quantified the similarity between latent states by calculating rank correlation coefficients between their respective frequency distribution of overlap patterns (see Methods and Materials). This analysis revealed highly concordant patterns of bias between each peak caller, with positive rank correlations between HMM states in 32/32 experiment sets (median 𝜌 = 0.9), 32/32 (median 𝜌 = 0.71) and 32/32 (median 𝜌 = 0.74) for MACS2 v GEM, MACS2 v SISSRs, and GEM v SISSRs, respectively.

### Regions of Bias Frequently Contain Strong Enrichment of TF Specific Motifs

A key question we sought to answer was whether biased regions – regions of poor reproducibility between replicates or experiments – contain low quality data. If regional bias is an indicator of low quality data, it is prudent to filter biased regions from ChIP-seq data. Alternatively, if biased regions contain valuable, high quality data then popular tools such as IDR and ChIP-R may be inappropriately filtering peaks in such regions through their consensus-based approaches.

We used the enrichment of known TF motifs^18^ as a proxy for data quality; we reasoned that if regions with greater bias are not strongly enriched for the binding motifs associated with their TFs, then this suggests a lack of evidence of genuine binding and justifies their exclusion. Within a set of regions belonging to the same latent state, we quantified the severity of bias as the percentage of peaks belonging to the most dominant experiment or replicate, in experiment sets and replicate sets, respectively.

We hypothesised that if regional bias is associated with poorer quality data, then states with less severe bias feature consistently higher motif rates. To test this, we performed pairwise comparisons between all recovered states, within experiment and replicate sets; we measured the probability that the least biased state in a pair contained the highest motif rates.

States with greater bias contained poorer enrichment of motifs in replicate sets but not experiment sets. The least biased state contained the greatest motif enrichment in only 54% (83/155) (p=0.428) of pairs of states in experiment sets and 60% (78/130) (p=0.036) in replicate sets. Further, the ratio of bias severity and the ratio of motif enrichment between pairs of states was negatively correlated in replicate sets (-0.398, p=1e-04), meaning that the more biased a state is towards a single replicate, the poorer its motif enrichment compared to other states. The log-log relationship between bias severity and motif enrichment was not statistically significant in experiment sets (𝜌=-0.207, p=0.134).

Subtle differences in sequence content of motif sites, or different propensities for non-canonical binding between regions may lead to slight differences in motif enrichment that do not reflect genuine differences in data quality. From this we reasoned that states with large decreases in motif enrichment relative to other regions are more likely to represent genuine differences in quality and would likely be of greater concern to researchers.

To reflect this in our analysis, we used a logistic Generalised Additive Model (GAM)^19^ to model the probability that the least biased state contained the greatest motif enrichment as a function of the absolute magnitude of the difference in motif enrichment between pairs of states.

Results were similar for replicate and experiment sets; the greater the absolute difference in motif enrichment between states, the greater the probability that the least motif enriched state is also the most biased state. To assess the statistical significance of these results, we used a permutation test (see Methods and Materials) to test the rank correlation between the absolute log ratio of motif enrichment between pairs of states and a binary indicator variable equal to 1 if the least biased state contained the greatest motif enrichment and 0 otherwise. The association was statistically significant for experiment sets (𝜌=0.226, p=0.013), but not for replicate sets (𝜌=0.107, p=0.271).

Despite the overall association between greater bias and poorer motif enrichment, our data revealed many exceptions to this trend. Beyond pairs of states, the most biased state overall contained the greatest motif enrichment in 39% (15/38) of experiment sets and 44% (27/62) of replicate sets. Across both experiment and replicate sets, over 40% of these high motif enrichment biased regions featured >70% of their peaks originating from a single experiment or replicate, meaning they are not merely regions of subtle disagreement between replicates or experiments. Thus, despite the broad statistical association between greater bias and poorer motif enrichment, filtering regions with inconsistent data between experiments and replicates may exclude valid TF binding sites.

### Regions of Poor Data Quality Feature Systematically Low ChIP-signal

We investigated whether ChIP-signal, defined as the fold-change in read count from control, could be used as a source of evidence of regional data quality that is independent of, yet complementary to motif analysis. We reasoned that if regions with lower motif rates consistently have lower ChIP-signal, it suggests that differences in motif rates do indeed reflect differences in underlying data quality. Further, if ChIP-signal can reliably predict motif rates, regional ChIP-signal may serve as a suitable metric for corroborating such evidence when discerning the aggregate data quality of a region.

To validate the connection between ChIP-signal and motif enrichment, we examined whether peaks that contained motifs featured higher ChIP-signal than peaks without motifs across each of our sampled experiment sets that targeted a TF with a motif in the JASPAR database^20^. We calculated the ratio of mean signal for peaks with and without a motif within a given replicate, and aggregated this result by taking the median across all replicates within a given experiment set. This analysis revealed that peaks containing motifs had greater ChIP-signal in 95% (38/48) of experiment sets, with peaks containing a motif featuring a median of 36% greater ChIP-signal than peaks without motifs. This result shows that ChIP-signal is indeed related to motif enrichment in the vast majority of experiment sets, validating its use as a complementary quality metric to motif enrichment.

To compare ChIP-signal across regions, we defined a “signal score” that measures the ratio of average fold-change values between different regions in the same experiment or replicate set. To control for differences in the fold-change distributions of different replicates, this score only compares the average fold-change of *same* replicate between *different* regions. Similarly, this score only uses comparisons between the same overlap pattern, e.g. the fold-changes of peaks belonging to “T T F F” unions within one region are only compared to the fold-changes of peaks forming “T T F F” unions within another. The signal score ratio is calculated as the geometric mean of the ratios of average fold-changes for each replicate within each overlap pattern between regions. A formal description of our approach can be found in Methods and Materials.

If signal score is to be used as a corroborating source of evidence to motif enrichment, states with higher motif enrichment should be consistently assigned higher signal score in between-state comparisons.

We uncovered a positive association between signal score and motif enrichment in pairwise state comparisons in experiment sets, with the state with the highest ChIP-signal also containing the highest motif enrichment in 66% (96/144) (p=0.001) of pairs. Curiously, regional ChIP-signal showed little association with motif enrichment in replicate sets overall (see Figure 4) as the state with the highest ChIP-signal also contained the highest motif enrichment in only 53% (63/117) (p=0.412) of pairs. This suggests that while ChIP-signal may be associated with motif enrichment between individual peaks, regional ChIP-signal is not a reliable predictor of regional motif enrichment in replicate sets.

**Figure 3.**
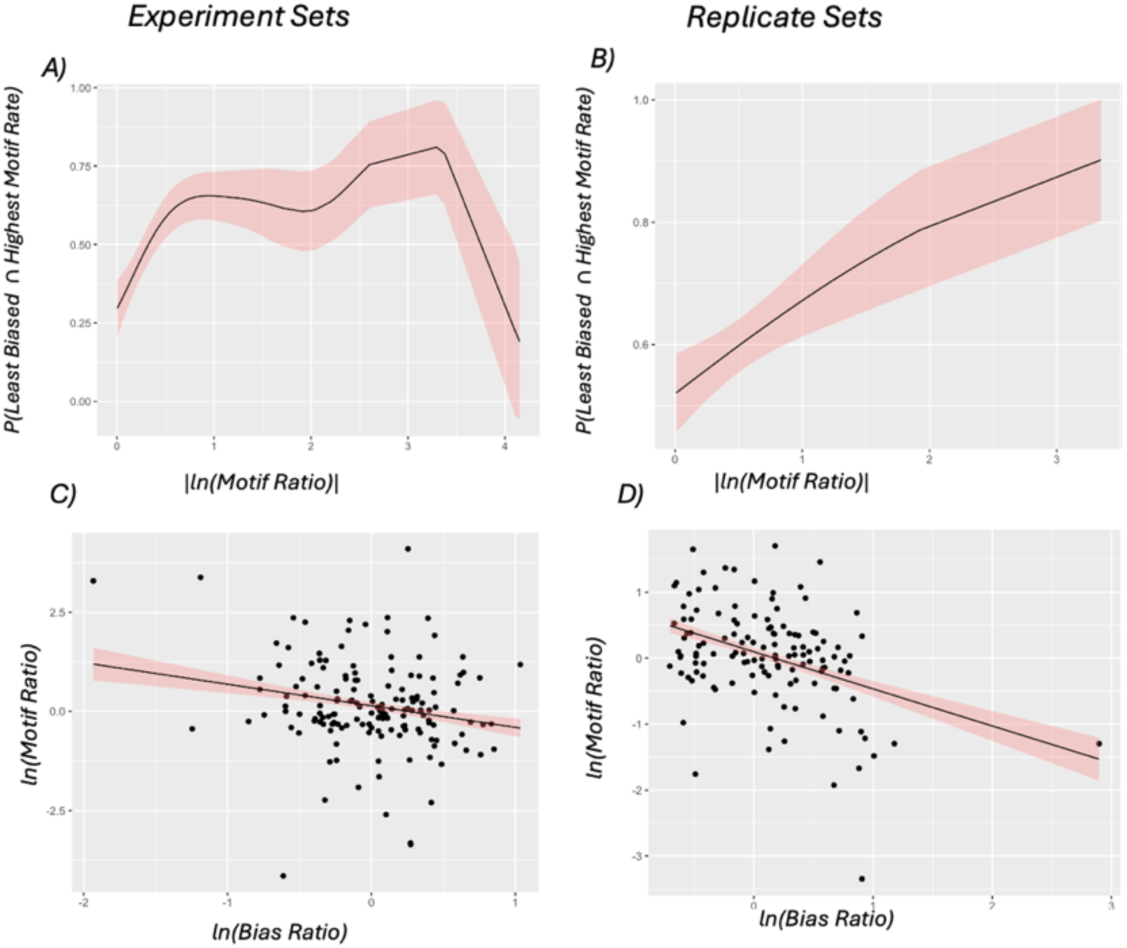
As the magnitude of the difference in motif enrichment increases between pairs of states, the least biased state becomes more likely to be the least motif enriched in Experiment Sets (A) And Replicate Sets (B): The probability that the more biased state in all pairs of states also has the lowest motif enrichment, expressed as a function of increasing magnitude of the log motif ratio between the two states for experiment sets and replicate sets, respectively. Smooth curves and 95% confidence intervals were estimated using a logistic General Additive Model implemented in the mgcv R-package. **C) & D): The proportional relationship between the difference in bias severity and the difference in motif enrichment between pairs of states in Experiment Sets C) and Replicate Sets D):** The relationship between the log ratio of bias severity and the log ratio of motif enrichment for pairwise state comparisons, within experiment sets and replicate sets, respectively. Regression lines were fit using linear regression.

**Figure 4:**
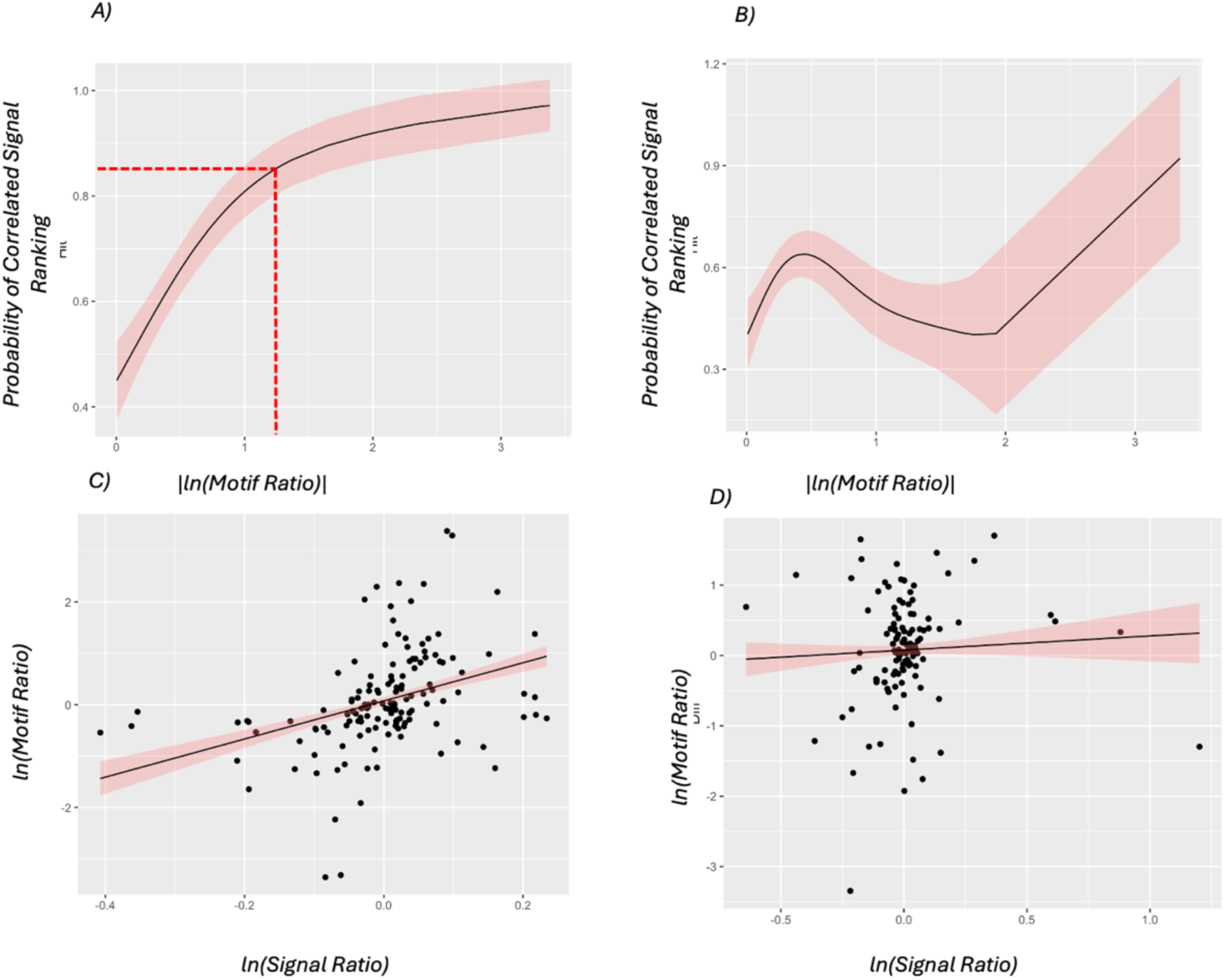
A & B) As the magnitude of the difference in motif enrichment increases between pairs of states, the state with the highest ChIP-signal becomes more likely to be the most motif enriched in Experiment Sets (A) but not Replicate Sets (B): The probability that the more biased state in a pairwise comparison also has the lowest motif enrichment, expressed as a function of increasing magnitude of the log motif ratio between paired states for experiment sets and replicate sets, resp. Dashed red lines indicate the magnitude of the log motif ratio beyond which ChIP-signal and motif rate ranks have a >85% chance of matching. Smooth curves and 95% confidence intervals were estimated using a logistic GAM**. C) & D):** The proportional relationship between the difference in ChIP-signal and the difference in motif enrichment between pairs of states in Experiment Sets C) and Replicate Sets D): The relationship between the log ratio of ChIP-signal and the log motif ratio for pairwise state comparisons within experiment sets and replicate sets, resp. Regression lines were fit using linear regression.

As larger differences in motif enrichment between states may be of greater concern to researchers than smaller differences, we also investigated whether signal score becomes more likely to select the most motif enriched state as the magnitude of differences in motif enrichment between states grows larger.

We used a logistic GAM to model the probability that the state with the greatest signal score also contained the greatest motif enrichment as a function of the absolute difference in motif enrichment between states. In experiment sets, the effectiveness of using signal scores to select the most motif enriched state improved substantially as the magnitude of differences in motif enrichment between states grew larger (𝜌=0.346, p=0.0001); states had a greater than 85% chance of having a higher signal rank than the state it was compared to if it also had greater than 3.3-fold higher motif rate in experiment sets (see Figure 4A). In replicate sets, the state with the greatest ChIP-signal was no more likely to select the state with the highest motif enrichment when the absolute difference in motif enrichment was large than when it was small (𝜌=0.032, p=0.297).

Overall, regional ChIP-signal is strongly associated with regional motif rates in experiment sets. We previously discovered that many regions of bias between experiments contained strong motif enrichment, showing that filtering peaks by consensus is not a reliable approach integrating multiple ChIP-seq experiments. Our results suggest that a combined ChIP-signal and motif analysis may be a more suitable method for distinguishing experiment-biased regions of high and low quality

### Regions of Less Accessible Chromatin are Hotspots for Inter-experiment Regional Bias

Prior research has demonstrated that ChIP-seq experiments extract relatively fewer reads from closed chromatin compared to regions of open chromatin due to the greater resistance to fragmentation featured by compact chromatin^21,22^. We hypothesised that technical variation in the sonication step between laboratories may lead to bias in regions of less accessible chromatin.

To test this hypothesis, we selected all experiment sets with at least one HMM state representing experiment bias (>70% of peaks unique to one experiment, see Methods and Materials). We defined genomic intervals by the coordinates of the first and last peak of each state segment identified by the Viterbi algorithm (see Methods and Material); we then quantified the ATAC-seq^23^ signal (reads per Mb) within the intervals belonging to each HMM state. We categorised a biased region as residing in less accessible chromatin if it featured at least four-fold fewer ATAC-seq reads per megabase than at least one unbiased region within the same experiment set. Finally, we then recalled peaks using MACS2, GEM and SISSRs and excluded experiment sets that did not yield a biased state across all three peak callers.

Our final filtered dataset contained 10 experiment sets, featuring 8 unique TFs across 3 human cell lines. We then used cell line specific RNA-seq^24^ to validate that that these experiment sets contained biased regions in compact chromatin. Peak proximal genes (<3Kb) in the biased regions of these experiment sets featured a median of 92.3% (ranging between 65% and 97%) decrease in expression levels compared to the genome wide average, confirming that these regions reside in less accessible chromatin.

Biased regions associated with less accessible chromatin occupied large (greater than 1 Mb) contiguous genomic regions that cleanly overlay visible troughs in ATAC-seq signal (Figure 5).

**Figure 5:**
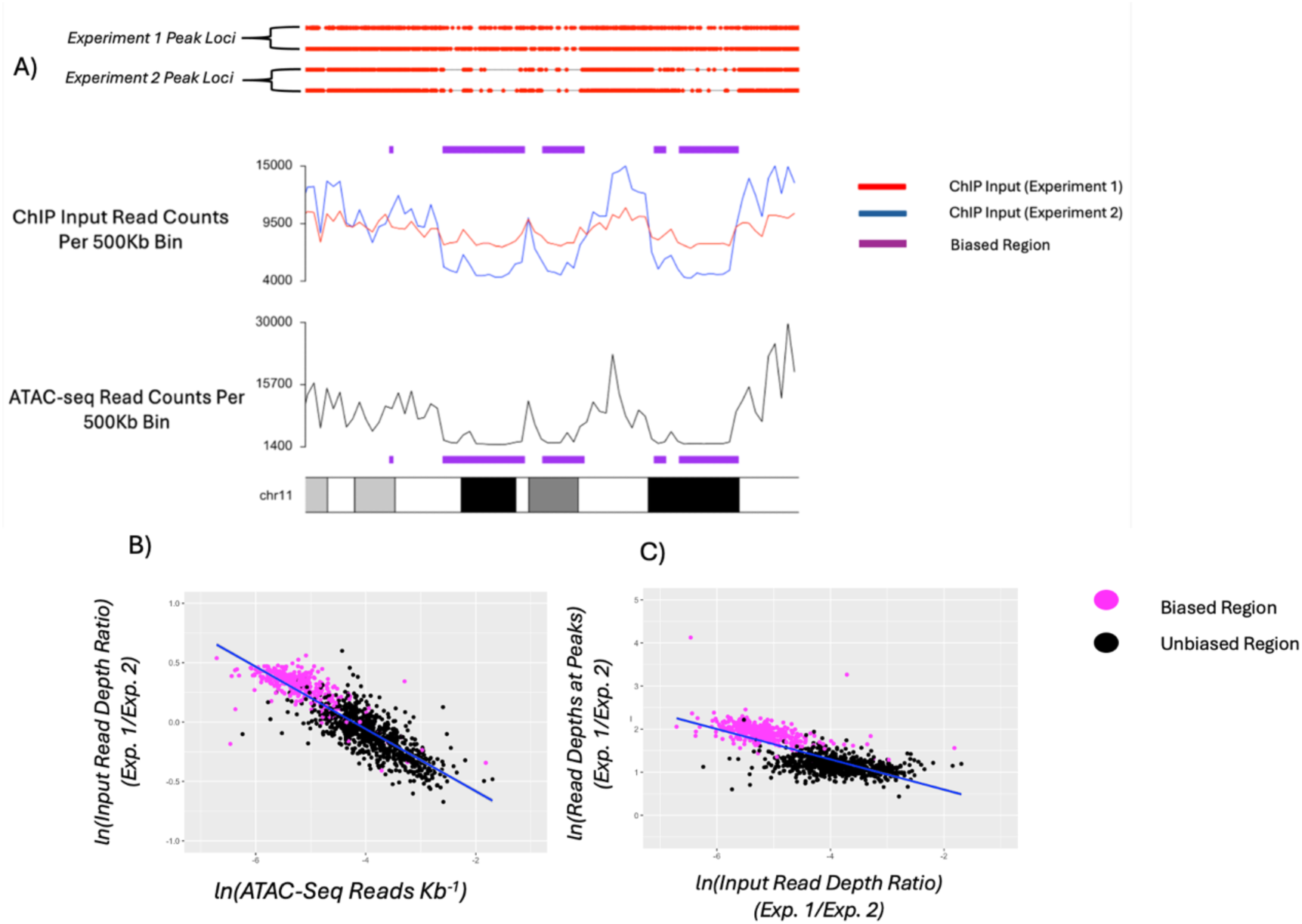
A) For regions of bias in less accessible chromatin, the experiment that discovers fewer peaks also features lower input read depth in closed chromatin: ATAC-seq, peak loci and ChIP input read counts for two experiments within a subset of chromosome 11 for experiment set CEBPG-K562. Purple rectangles mark the location of regional biases between these two experiments. Experiment 2 contains both fewer peaks and fewer input reads in biased regions than Experiment 1, suggesting that read extraction was more sensitive to chromatin state in Experiment 2. **B) For Experiment sets featuring Bias in Closed Chromatin, The ratio of ChIP-Input reads between Experiments is strongly associated with ATAC-seq read depth across the entire genome:** log-log plots of the ChIP Input ratio (Experiment 1/Experiment 2) and ATAC-seq reads across all contiguous genomic intervals assigned to an HMM state for CEBPG-K562. Regions annotated to biased states are highlighted purple. As DNA accessibility increases, the bias in background signal recovery decreases. **C) The ratio of ChIP-input reads is strongly predictive of the ratio of Read Depths at peaks between Experiments:** log-log plots of the ChIP Input ratio (Experiment 1/Experiment 2) and the ratio of read depths at peaks (Experiment 1/Experiment 2) across all contiguous genomic intervals assigned to an HMM state for CEBPG-K562, respectively. Experiment biased states are highlighted purple. The degree of bias in the number of peaks recovered between experiments is predicted by the degree of bias in the recovery of input reads. The experiment CEBPG-K562 consisted of three experiments, the comparison displayed here is between experiments ENCSR490LWA and ENCSR620VIC.

Strikingly, compact chromatin-biased regions featured the highest motif enrichment out of any other regions in 9/10 experiment sets with known binding motifs, featuring 23%-295% higher motif rates than the average of other states. To corroborate motif evidence, we compared the signal strength of peaks in less accessible chromatin using the signal score. In every experiment set, biased regions in less accessible chromatin featured comparable or higher signal strength compared to any other region (see Supplementary Table 1). The combination of strong motif enrichment and strong ChIP-signal presents strong evidence that for at least 9/10 experiment sets, peaks in less accessible chromatin represent genuine binding sites.

Of the eight unique TFs in our sample, only one (FOXA2) is broadly recognised as playing a functional role in closed chromatin^12^. Despite not being considered canonical pioneer factors, all TFs identified had between 7%-85% of their binding sites in less accessible chromatin regions, matching or exceeding that of FOXA2 (10%). This suggests that these TFs play previously unrecognised roles in compact or closed chromatin.

### Bias in Less Accessible Chromatin Arises Due to Technical, not Biological Variation Between Experiments

Biased regions we discovered in less accessible chromatin are composed of one experiment containing relatively abundant peaks (henceforth C.C abundant), and one or more experiments containing relatively sparse peaks (henceforth C.C sparse). It is well known that compact chromatin generates fewer reads in ChIP assays^21,22^, leading us to hypothesise that C.C abundant experiments were better able to extract reads than C.C sparse experiments, leading to much greater discovery of peaks in less accessible chromatin by C.C abundant experiments.

To test this hypothesis, we compared the read depth in less accessible chromatin in the input controls of C.C abundant and C.C sparse experiments, reasoning that if C.C sparse experiments are less able to extract reads from compact chromatin, this would be reflected in greater sensitivity to chromatin state in their ChIP input replicates. Read depth in C.C sparse experiments showed on average far greater sensitivity to chromatin state than C.C abundant experiments, with the log ratio of C.C abundant/sparse input read depth increasing linearly with decreasing chromatin accessibility in 7/10 experiment sets (see Figure 5A, Figure 5B, Supplementary Table 1). The relationship between chromatin state and input read depth disparity between experiments was powerful, with ATAC-seq read depth explaining a median of 50% of the variation in log read depth ratio between experiments genome wide (see Supplementary Table 1).

Finally, we tested the relationship between chromatin accessibility and the ratio of read depth in peaks for C.C abundant/sparse across the genome. This analysis revealed that C.C abundant experiments featured progressively higher read depths in peaks relative to C.C sparse experiments as chromatin accessibility decreased genome wide (see Supplementary Table 1). The fact that the same differences in sensitivity to chromatin state between experiments is observed in both treatment and input ChIP replicates shows that it is related to basic technical differences in the ChIP assay between laboratories.

### Biased Regions often Overlap With Structural Variants

We also investigated whether other regional features, such as GC content^25^ or copy number variation (CNV)^26^ correlated with bias between experiments. To investigate the influence of CNVs on regional bias, we took CNV annotations from the UCSC Genome Browser^27^ for the cancer cell lines K562 and HepG2 and tested whether peaks annotated to different states differed in their likelihood of overlapping different CNVs (either amplifications or deletions). The strong autocorrelation between adjacent peak annotations violates the assumptions of standard statistical tests, so we performed permutation tests by shuffling the order of Viterbi state segments and regional CNV annotations (see Methods and Materials). This analysis revealed that 52% (32/61) of experiment sets and 29% (26/91) of replicate sets featured statistically significant differences in CNV overlap between states. On average, the cell line K562 was more likely to feature statistically significant differences in CNV overlap between states in both experiment sets (𝜒^2^ =5.0941, *p*=0.024) and replicate sets (𝜒^2^ =3.097, *p*=0.0785) than the HepG2 cell line. The median percentage of peaks for the most biased state within each CNV is detailed in Supplementary Table 2.

We further used linear regression to test if the proportion of overlap with different CNVs (amplifications or deletions) and GC content was associated with bias severity and motif enrichment between pairs of states in experiment and replicate sets. Because the same HMM state may appear in multiple pairwise comparisons, we used permutation tests by permuting the state labels within experiment and replicate sets (see Methods and Materials). This analysis revealed little association between overlap with motif enrichment and bias severity for both CNV overlap and GC content (Supplementary Table 3).

### Standard Quality Control Metrics Do Not Predict the Occurrence or Severity of Regional Bias

We investigated whether standard ChIP QC metrics could predict either the occurrence or severity of biased regions between experiments or replicates. We used the R package ChIPQC^28^ to quantify the Relative Strand Cross Correlation, the fraction of reads in peaks, the uniformity read enrichment, and the total read depth for each replicate in our sample of 90 experiment sets^29^. Across each quality metric, we used logistic regression to test the association between the minimum score achieved across each replicate and the probability of regional bias being detected in that replicates respective experiment or replicate set. We then used linear regression for each quality metric to test the association between the minimum score across each replicate and the ratio of the maximum/mean bias severity observed within each experiment and replicate set. This analysis little association between standard ChIP QC metrics and regional bias, and no analyses were statistically significant after adjusting for multiple comparisons (see Supplementary Table 4).

## DISCUSSION

Regional bias is a widespread issue driving systematic differences between ChIP-seq replicates and experiments. Strikingly, biased regions often contain clear evidence of containing genuine TF binding, despite not being reproduced between matching experiments. Standard methodology for integrating ChIP-seq replicates and experiments such as IDR and ChIP-R cannot account for this phenomenon, and so may unknowingly filter valuable data via the emphasis on consensus peaks.

Inter-experiment bias can lead to dramatic differences in the biological information provided by different experiments, as evidenced by the prevalent bias in regions of less accessible chromatin. We provide evidence that eight unique TFs have the capacity to bind to closed chromatin, and that this fundamental aspect of their biology was starkly absent from some experiments but not others. Closed chromatin binding is not considered a defining feature of the majority of TFs in our sample, yet they feature binding sites in less accessible chromatin at rates matching or exceeding that of FOXA2, a well-recognised pioneer factor. This suggests that the empirically variable ability (amongst contributors to ENCODE) to reliably detect binding in regions of less accessible chromatin may have led to an underestimate of the number of TFs that exhibit this capacity. Lending favour to this hypothesis is the fact that our analysis was limited to cell lines with available ATAC-seq data, leaving the possibility that even within the curated ENCODE datasets there remain TFs with undiscovered activity in closed chromatin.

The discovery of bias in less accessible chromatin also points towards certain technical factors as contributors to bias; our data suggest that the fragmentation step in the ChIP-seq may lead to inter-experiment bias in regions of less accessible chromatin. We showed that the observed regional bias resulted from differing baseline read counts in less accessible chromatin across laboratories, a variable for which DNA fragmentation (primarily via sonication) plays the most salient role^21,22^. It has long been known that closed chromatin is resistant to sonication and thus underrepresented in ChIP DNA libraries, but our results are the first to provide evidence that this phenomenon can vary between experiments, leading to dramatic effects on reproducibility.

The technical sources of other biased regions remain entirely unknown but the ubiquity of this issue across thousands of datasets suggests that in general, technical variation tends to produce region specific effects. Improving the reproducibility of ChIP-seq will require a careful review of the existing technical standards, with further research required to identify the technical sources of bias.

The computational methodology we present in this paper shows great potential for ameliorating some the worst impacts of bias. We have provided methodology capable of efficiently identifying systematic bias between ChIP-seq replicates, and at least for inter-experiment comparisons, A combination of motif and signal analysis can determine whether regions of bias should be discarded or integrated into a joint set of peaks. The importance of this latter point is underscored by the variable discovery of binding in closed chromatin; rescuing the biological information within some biased regions is vital, but it is equally important to prevent datasets from becoming populated with regional artifacts.

Encouragingly, the metrics we used to define the quality of experiment biased regions rest upon clear biological principles; our data suggest the strongest indicator of quality is the relative enrichment of a given TFs known sequence specificities. This statement is supported by the strong association between motif rates and ChIP-signal, where regions with markedly lower motif rates (3.3 fold for experiment sets) also contained the weakest binding signal in greater than 85% of cases. We employed this approach effectively in our analysis of less accessible chromatin, where a combined motif and signal analysis confirmed the validity of binding sites in less accessible chromatin for at least 9/10 experiment sets and 7/8 unique TFs.

Regions of bias between intra-experiment replicates were not amenable to the combined motif and signal analysis. ChIP-signal strength and motif enrichment were uncorrelated within replicate biased regions, indicating that either the enrichment of known motifs, the strength of ChIP-signal - or both – are not consistent markers of data quality in replicate biased regions. More research is needed to understand the characteristics of replicate biased regions, and why they differ from regions of bias between experiments. In the meantime, researchers must treat replicate biased regions with caution and closely monitor the influence their inclusion or exclusion has on the results of their analysis.

Despite these preliminary successes, many challenges remain for the analysis of ChIP-seq data considering this newfound bias. In our signal and motif analysis, subtler differences in motif enrichment between states were not consistently correlated with differences in signal score. It is possible that small differences in motif enrichment simply reflects slightly different sequence affinities between regions. Alternatively, it may be that the signal score is not sufficiently precise to represent differences in data quality when those differences are small.

More research is needed into the features that separate biased regions of high and low quality so that their impact on ChIP-seq data is minimised. In the meantime, our work provides researchers with the means to detect regional bias in their own data, and the ability to remove or retain biased regions that are of distinctly low or high quality.

## METHODS AND MATERIALS

### Hidden Markov Model Finds Systematic Regional Bias

The peak union sequence is a sequence of unified intervals sorted by genomic locus and formed by merging all overlapping peaks across each replicate in an experiment set.

With this construction, the interval spanned by any peak in any replicate is a subset of the interval of exactly one peak union. For peaks that are found in only one replicate, the “union” representing this peak is simply the peak itself. Each peak union is assigned a T/F tuple where each “T” and “F” denote whether a peak from each replicate is present or missing from the interval spanned by the peak union. As a convention, replicates from the same experiment are adjacent in the T/F tuple e.g. “T T F F” refers to a peak identified in both replicates of the first experiment, but neither replicate from the second. This encoding allows for systematic regional variation to be detectable as changes in the distribution of T/F tuples across the peak union sequence.

HMMs^11^ are employed to identify changes in the distribution of T/F tuples across the peak union sequence, and cluster the sequence into segments of discrete, latent “states”. The BIC^13^ was used to select state number selection. The Baum-Welch algorithm^29^ was used to determine HMM parameters. The Viterbi algorithm^30^ was used to annotate the peak union sequence with the most likely state sequence, which in turn assigned a state to each genomic locus occupied by a peak. All analyses were performed using the R package hmm.discnp (available in CRAN).

### Defining State of Bias

The distribution of overlap combinations for each state was used to identify states that represent bias towards a specific replicate or experiment. A state was categorised as “experiment biased” or “replicate biased” if over 70% of its overlap combinations originated uniquely from the same experiment or replicate respectively. We also use a continuous definition of bias, referred to as “bias severity”, which is quantified by the percentage of peaks within a region originating uniquely from the same experiment or replicate.

### Defining Contiguous Regions of Bias

The Viterbi sequence consists of segments of reoccurring states. We use these segments to identify the contiguous genomic region occupied by each segment, and therefore the set of contiguous regions occupied by each HMM state. The start and end coordinates of each contiguous region is defined by the coordinates of the first and last peak union in the segment respectively.

### ENCODE Data Curation and Analysis

To quantify the frequency of systematic regional bias in ChIP-seq data, we ran HMMs across a large, curated portion of human ENCODE ChIP-seq data. For regional bias across experiments, we paired all datasets investigating the same transcription factor in the same tissue, forming matched experiments termed “experiment sets”. For all ENCODE experiments that could not be matched, replicates from these experiments formed “replicate sets”, and were used to investigate regional bias between replicates of the same experiment. To avoid potential biological confounders, we filtered experiment/replicate sets to only those that contain data with no additional chemical treatments. We also filtered experiments containing fewer than 1000 total peaks, ensuring any detected bias was not the result of anomalously sparse peak numbers.

The sample curation results in a total of 229 experiment sets and 1792 replicate sets. For each experiment/replicate set, peak union sequences were constructed as described above using IDR pseudo-replicated peaks (see ENCODE Terms and Definitions at https://www.encodeproject.org/data-standards/terms/).

HMMs were run as described above for all peak union sequences in both datasets. The BIC was used to select the optimal number of states, and an experiment or replicate set was considered to contain systematic bias if its optimal state number was greater than 1.

The BIC has previously been shown to be provide consistent estimate the number of latent states in HMMs^31^, meaning the BIC is increasingly likely to select the correct number of states as the length of the sequence the HMM is trained on increases.

To validate the accuracy of the BIC to detect bias directly, we used randomised sequences to measure the frequency with which the BIC erroneously detects bias in random noise. Truly random sequences were constructed by shuffling the order of overlap combinations within each peak union sequence. HMMs were run on the randomised sequences in order to empirically estimate the rate of truly random sequences identified as biased (type I error rate).

### Comparing Bias across Peak Callers

We investigated regional bias across three peak callers: GEM^16^ (version 3.4), MACS2^15^ (version 2.2.7.1) and SISSRs^17^ (version 1.4) These peak callers were selected because they each employ distinct methods for the identification of candidate peaks and subsequent statistical filtering. Further, each peak caller has been found to perform comparably in benchmark studies^32,33^, making each tool a reasonable representation of a high-quality analysis that researchers in the field may employ.

A random sample of 90 experiment sets was taken, and peaks were called for each replicate by each peak caller using default parameters. For MACS2 and GEM, all control files in the ENCODE portal were merged for peak calling. SISSRs did not accept multiple controls, so the control replicate with the highest read depth was chosen for peak calling. HMMs were trained on peak union sequences for each peak caller separately, and the BIC was used to select the optimal number of states. As done previously, an experiment set was defined as containing regional bias if the BIC selected an optimal state number >1.

To compare biased regions between peak callers we defined a common set of peak loci by taking the intersection of peak union sequences derived from peaks called by each peak caller. This results in a set of peak loci that are unanimously called by each peak caller, but do not necessarily share the same overlap pattern across replicates.

Viterbi annotations were compared between the (peak caller specific) HMMs to pair states for comparison. States were paired between HMMs such that each pair contained the greatest number of overlapping Viterbi annotations.

We quantified the similarity between HMMs trained on different peak callers by calculating the rank correlation coefficients between the distribution of overlap patterns of paired states and then taking the median of these coefficients across all pairs of states in the comparison.

### Motif Analysis

Rates of motif occurrences across regions were performed using FIMO version 5.5.5^34^. Motif profiles for each transcription factor were sourced from JASPAR 2022^20^ redundant collection. All motifs associated with each transcription factor were combined into “motif sets” for scanning with FIMO. A peak union was considered to contain a motif if FIMO returned at least one significant match for at least one member of the relevant motif set.

### Regional ChIP-Signal

For a set of *k* replicates, let **P** be the set of 2*^k^*-1 unique binary tuples representing each possible overlap pattern, where for all *pi* ∈ **P**, 𝑝_𝑖j_ is equal to 1 if the *j*th replicate contributes a peak to the overlap pattern corresponding to *pi*. Further, let

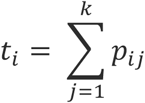

Represent the total number of replicates contributing to the *i*th overlap pattern. Finally, let 𝑆_𝑖j_ be the set signal values belonging to peaks occurring in an instance of the *i*th overlap pattern in state 𝑆 belonging to the *jth* replicate, and 𝑁_𝑖j_ the total number of such observations.

We then define the replicate-overlap pattern specific score for state 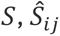 as:

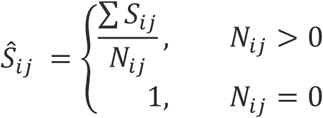

And the overall signal score for state 𝑆, 𝑆𝑐 is calculated as:

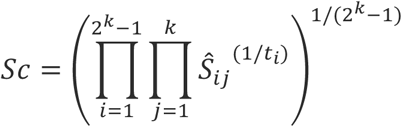

In practice, some overlap patterns will not feature enough observations in all states. In this paper, only overlap patterns with over 40 observations in each state were used for the calculation of signal scores.

### Permutation Tests for Pairwise State Comparisons

Multiple analyses throughout this paper rely on performing pairwise comparisons between states within experiment or replicate sets to test associations between regional characteristics. In these analyses, a given HMM state can appear in multiple pairwise comparisons, and therefore the assumption of independent observations, essential for standard statistical tests, is violated. We account for this by performing permutation tests where the state level variables are randomly reshuffled within each experiment or replicate set before sorting states into pairs. This approach enables us to test the null hypothesis of no association between the variables in question whilst maintaining the non-independence induced by the same state appearing in multiple comparisons. For each test, we perform n=10,000 permutations and record the value of the relevant test statistic in each iteration. Empirical p-values are then derived by measuring the proportion of randomly derived test statistics that are greater than the test statistic observed in the non-randomised data.

### Permutation Tests for CNV Overlap

We investigated the association between regional bias and Copy Number Variation (CNV) in the K562 and HepG2 cell lines. We downloaded copy number variation annotations from the UCSC Genome Browser^24^ for cell lines K562, and HepG2 (https://genome.ucsc.edu/cgi-bin/hgTrackUi?db=hg18Cg=wgEncodeHudsonalphaCnv), and lifted genomic coordinates from genome assemblies hg18 to hg38 using the UCSC LiftOver tool^26^. To measure the association between states and CNVs, for each replicate and experiment set we calculated the proportion of peaks within each state that fell into a region annotated to one of two CNV categories (amplification or deletion) or a region annotated as not containing a CNV. Due to the strong autocorrelation between adjacent state annotations in the Viterbi sequence, we tested these associations for statistical significance with permutation tests; state annotations were permuted in blocks to defined by contiguous state segments (see above) and CNV annotations were also randomly permuted. This approach allowed us to test the association between patterns of bias and CNVs whilst keeping the spatial distribution of peak unions and the statistical dependency between adjacent peaks fixed. For these tests we used n=10000 permutations, and derived p-values by calculating the proportion of test statistic values (𝜒^2^) in our empirical null distribution that exceeded the 𝜒^2^ observed in the real data.

### Permutation Tests for Regional Bias Hotspots

We investigated whether some genomic regions were more prone to regional bias than others, and whether the propensity to feature regional bias differed between cytoband categories. The human genome assembly GRCh38 was divided into 100Kb bins, and for each bin, the proportion of overlapping states that were the maximum biased states in their respective experiment sets was calculated. We used an F statistic to measure the overall association between Giemsa staining band categories and the odds that a state overlapping each category was the most biased state in its experiment set. To account for the non-independence between adjacent cytoband labels, assessed the statistical significance of this F statistic by performing 10,000 random permutations of each block of recurring, identical cytoband labels. For each permutation we recorded the magnitude of the F statistic for each permutation.

### Defining Less Accessible Chromatin

To identify potential instances of bias in less accessible chromatin, a set of candidate experiment sets was constructed from ENCODE called peaks, where experiment sets with cell line matching ATAC-seq data and least one experiment biased state (defined above) was retained. Experiment sets were further filtered to those that contained at least two experiments that do not feature an ENCODE Audit failure. Experiments with an audit failure were not filtered so long as they formed part of an experiment set with two or more experiments without an audit failure. The hypothesis we investigated pertained to inter-lab differences in sensitivity to chromatin state, so experiment sets that did not feature experiments from at least two laboratories were excluded. To assist interpretability and avoid redundancy, duplicate experiments from the same laboratory were removed from each experiment set by removing the experiment which contained the fewest peaks.

Experiment biased states were then considered to be located in less accessible chromatin if the contiguous regions they occupied contained at least 4-fold fewer ATAC-seq reads per megabase than at least one “unbiased” state in the experiment set.

We then re-called peaks using GEM^16^ MACS2^15^ and SISSRs^17^, and excluded experiment sets that did not include regions of bias across all three peak callers.

We further validated this analysis by quantifying the absolute gene expression levels of peak-proximal (within 3kb) genes in regions of less accessible chromatin; if these regions are indeed less accessible, they should display decreased gene expression.

We compared the relationship between chromatin accessibility and ChIP-seq read depth (both input and treatment) between experiments with relatively abundant peaks in closed chromatin (C.C abundant) and experiments with relatively sparse peak counts in closed chromatin (C.C sparse). Read densities must first be normalised to account for replicate level differences in read abundance, however, normalising replicates by total read counts would confound results if the contribution of reads in less accessible chromatin to total number of reads differed substantially between replicates. To account for this, we used linear regression to measure the difference in sensitivity to chromatin state between experiments by modelling the log ratio in ChIP-input read depth between experiments as a linear function of log ATAC-seq read depth. The slope of this linear relationship reveals whether the input-read depth ratio changes as a function of chromatin accessibility, and therefore which, if any experiment shows greater sensitivity to chromatin state. Multiplicative constants become intercept terms on the log scale and do not obscure the relationships (slopes) between variables; no normalisation was necessary for this analysis.

### Copy Number Variation and GC Content

We downloaded copy number variation annotations from the UCSC Genome Browser^27^ for cell lines K562, GM12878 and HepG2 (https://genome.ucsc.edu/cgi-bin/hgTrackUi?db=hg18Cg=wgEncodeHudsonalphaCnv), and lifted genomic coordinates from genome assemblies hg18 to hg38 using the UCSC LiftOver tool^27^. We quantified the proportion of peaks overlapping with insertions and deletions, and measured the correlation between CNV overlap and bias severity, and CNV overlap and motif enrichment between states.

We also compared the GC content of peaks between states, measured the correlation between GC content and bias severity, and GC content and motif enrichment between regions.

### QC Analysis

We investigated whether regional bias between replicates or experiments could be predicted by standard ChIP-seq quality metrics. We used the R package ChIPQC^28^ to calculate the Relative Strand Cross Correlation Coefficient^28^, Sum of Squared Differences^28^, Fraction of Reads in Peaks^28^, and the total read depth for the MACS2 called peaks of our 90 sampled experiment sets. One experiment was excluded from this analysis after ChIPQC failed to run on this dataset. For each quality metric, we used logistic regression to investigate the association between the probability of regional bias occurring within a given experiment or replicate set, and the minimum quality score observed for an experiment or replicate within that set. Further, for each quality metric, we used linear regression to measure the association between the severity of bias within a given experiment set (defined as the maximum bias severity of any state divided by the average of all states), and the minimum quality score observed across any experiment or replicate within that set. P values were corrected using the Holm-Bonferroni correction^35^.

## CODE AND DATA AVAILABILITY

Code for constructing the peak union sequence, bias discovery with HMMs, and signal score calculation is implemented in the R package ChIPRegions, available on github https://github.com/Olly98/ChIPRegions. Scripts for the ENCODE wide bias analysis, and bias quantification for MACS2, GEMR and SISSRs called peaks are also available in this code repository, with the results for these analyses available on Zenodo DOI:**10.5281/zenodo.18409232**. Peaks called with MACS2, GEMR and SISSRs are available on Zenodo DOI:**10.5281/zenodo.1840G232**.

## ACKNOWLEDGMENTS

The project received no external funding.

## References

1. Mundade, R., Ozer, H. G., Wei, H., Prabhu, L. & Lu, T. Role of ChIP-seq in the discovery of transcription factor binding sites, differential gene regulation mechanism, epigenetic marks and beyond. Cell Cycle 13, 2847–2852 (2014).

2. ENCODE Project Consortium. An integrated encyclopedia of DNA elements in the human genome. Nature **489**, 57–74 (2012).

3. Li, Q., Brown, J. B., Huang, H. & Bickel, P. J. Measuring reproducibility of high-throughput experiments. Ann. Appl. Stat. 5, (2011).

4. Jalili, V., Cremona, M. A. & Palluzzi, F. Rescuing biologically relevant consensus regions across replicated samples. BMC Bioinformatics 24, 240 (2023).

5. Yang, Y. et al. Leveraging biological replicates to improve analysis in ChIP-seq experiments. Comput Struct Biotechnol J **9**, e201401002 (2014).

6. Czipa, E. et al. ChIPSummitDB: a ChIP-seq-based database of human transcription factor binding sites and the topological arrangements

7. Newell, R. et al. ChIP-R: Assembling reproducible sets of ChIP-seq and ATAC-seq peaks from multiple replicates. Genomics 113, 1855–1866 (2021).

8. Amemiya, H. M., Kundaje, A. & Boyle, A. P. The ENCODE Blacklist: Identification of Problematic Regions of the Genome. Scientific Reports **9**, 9354 (2019).

9. Yu, Y., Mai, Y., Zheng, Y. & Shi, L. Assessing and mitigating batch effects in large-scale omics studies. Genome Biology 25, 254 (2024).

10. Sprang, M., Andrade-Navarro, M. A. & Fontaine, J.-F. Batch effect detection and correction in RNA-seq data using machine-learning-based automated assessment of quality. BMC Bioinformatics 23, 279 (2022).

11. Rabiner, L. & Juang, B. An introduction to hidden Markov models. IEEE ASSP Magazine 3, 4–16 (1986).

12. Liu, N. et al. FOXA1 and FOXA2: the regulatory mechanisms and therapeutic implications in cancer. Cell Death Discovery 10, 172 (2024).

13. Neath, A. A. & Cavanaugh, J. E. The Bayesian information criterion: background, derivation, and applications. WIREs Computational Statistics 4, 199–203 (2012).

14. Barcia, J. J. The Giemsa Stain: Its History and Applications. Int J Surg Pathol 15, 292–296 (2007).

15. Zhang, Y. et al. Model-based Analysis of ChIP-Seq (MACS). Genome Biol **9**, R137 (2008).

16. Guo, Y., Mahony, S. & Gifford, D. K. High Resolution Genome Wide Binding Event Finding and Motif Discovery Reveals Transcription Factor Spatial Binding Constraints. PLoS Comput Biol 8, e1002638 (2012).

17. Narlikar, L. & Jothi, R. ChIP-Seq Data Analysis: Identification of Protein–DNA Binding Sites with SISSRs Peak-Finder. in Next Generation Microarray Bioinformatics (eds. Wang, J., Tan, A. C. & Tian, T.) vol. 802 305–322 (Humana Press, Totowa, NJ, 2012)

18. Boeva, V. Analysis of Genomic Sequence Motifs for Deciphering Transcription Factor Binding and Transcriptional Regulation in Eukaryotic Cells. Front. Genet. 7, (2016).

19. Wood, S. N., Goude, Y. & Shaw, S. Generalized Additive Models for Large Data Sets. Journal of the Royal Statistical Society Series C: Applied Statistics 64, 139–155 (2015).

20. Castro-Mondragon, J. A. et al. JASPAR 2022: the 9th release of the open-access database of transcription factor binding profiles. Nucleic Acids Res 50, D165–D173 (2022).

21. Auerbach, R. K. et al. Mapping accessible chromatin regions using Sono-Seq. Proc Natl Acad Sci U S A 106, 14926–14931 (2009).

22. Teytelman, L. et al. Impact of Chromatin Structures on DNA Processing for Genomic Analyses. PLOS ONE 4, e6700 (2009).

23. Chromatin accessibility profiling by ATAC-seq | Nature Protocols. https://www.nature.com/articles/s41596-022-00692-9.

24. Wang, Z., Gerstein, M. & Snyder, M. RNA-Seq: a revolutionary tool for transcriptomics. Nat Rev Genet 10, 57–63 (2009).

25. Teng, M. & Irizarry, R. A. Accounting for GC-content bias reduces systematic errors and batch effects in ChIP-seq data. Genome Res. 27, 1930–1938 (2017).

26. Su, D., Peters, M., Soltys, V. & Chan, Y. F. Copy number normalization distinguishes differential signals driven by copy number differences in ATAC-seq and ChIP-seq. BMC Genomics 26, 306 (2025).

27. Casper, J. et al. The UCSC Genome Browser database: 2026 update. Nucleic Acids Research 54, D1331–D1335 (2026).

28. Carroll, T. S., Liang, Z., Salama, R., Stark, R. & De Santiago, I. Impact of artifact removal on ChIP quality metrics in ChIP-seq and ChIP-exo data. Front. Genet. 5, (2014).

29. Yang, F., Balakrishnan, S. & Wainwright, M. J. Statistical and computational guarantees for the Baum-Welch algorithm. in 2015 *53rd Annual Allerton Conference on Communication*, Control, and Computing (Allerton) 658–665 (2015). doi:10.1109/ALLERTON.2015.7447067.

30. Lou, H.-L. Implementing the Viterbi algorithm. IEEE Signal Processing Magazine 12, 42–52 (1995).

31. Boucheron S, Gassiat E (2007) An information-theoretic perspective on order estimation. In: Cappé, Olivier and Moulines, Eric and Rydén, Tobias, (eds) Inference in hidden Markov models, pp 565–601

32. Jeon, H., Lee, H., Kang, B., Jang, I. & Roh, T.-Y. Comparative analysis of commonly used peak calling programs for ChIP-Seq analysis. Genomics Inform 18, e42 (2020).

33. Thomas, R., Thomas, S., Holloway, A. K. & Pollard, K. S. Features that define the best ChIP-seq peak calling algorithms. Brief Bioinform bbw035 (2016) doi:10.1093/bib/bbw035.

34. Grant, C. E., Bailey, T. L. & Noble, W. S. FIMO: scanning for occurrences of a given motif. Bioinformatics 27, 1017–1018 (2011).

35. Holm, S. A Simple Sequentially Rejective Multiple Test Procedure. Scand J Statist.

