## Supplementary Tables for "Systematic Regional Bias is Widespread in ChIP-seq"

| **Targeted TF** | **Cell Line** | **ATAC vs Input ChIP Ratio Slope** | **ATAC vs Input ChIP Ratio R**^2^ | **ATAC vs Treatment ChIP Ratio Slope (C.C Sparse/C.C Abundant)** | **% Motif Increase** | **% ChIP-Signal Score Increase** | **Gene Expression**  **Decrease in Biased Regions** | **Experiment Accessions** |
| --- | --- | --- | --- | --- | --- | --- | --- | --- |
| ATF2 | GM12878 | 0.18 | 0.41 | 0.86 | 295% | 10% | -92% | ENCSR000BQK, ENCSR961PPA |
| ATF2 | HepG2 | -0.04 | 0.02 | 0.95 | 266% | 5% | -96% | ENCSR047BUZ, ENCSR908HWZ |
| ATF3 | K562 | -0.04 | 0.01 | 0.2 | 153% | 6% | -92% | ENCSR000DOG, ENCSR000BNU ,ENCSR028UIU |
| CEBPG | K562 | 0.25 | 0.68 | 0.53 | 60% | 15% | -89% | ENCSR490LWA,ENCSR620VIC, ENCSR058DRG |
| CREB1 | HepG2 | 0.22 | 0.26 | 0.67 | -10% | 7% | -97% | ENCSR331ORD, ENCSR112ALD |
| FOXA2 | HepG2 | -0.03 | 0.01 | 0.49 | 215% | 5% | -93% | ENCSR066EBK, ENCSR490AMH |
| JUN | HepG2 | 0.15 | 0.5 | 0.84 | 158% | 3% | -97% | ENCSR747VUU, ENCSR000EEK |
| JUND | HepG2 | 0.18 | 0.56 | 0.65 | 120% | 0% | -97% | ENCSR000BGK, ENCSR000EEI |
| NFE2 | K562 | 0.13 | 0.38 | 0.36 | 23% | 14% | -65% | ENCSR552YGL, ENCSR000FCC, ENCSR000FAF |
| ATF2 | K562 | 0.43 | 0.84 | 0.27 | 21% | 4% | -85% | ENCSR014ARU, ENCSR869IUD |

**Supplementary Table 1: Table of results for all 10 experiment sets identified as containing bias between experiments in less accessible chromatin:**

“ATAC vs Input ChIP Ratio” and “ATAC vs Treatment ChIP Ratio” refers to the relationship between log transformed ATAC-seq reads per Mb and log transformed ratio of input ChIP read depth (C.C sparse/C.C abundant, see Methods and Materials) and the log transformed ratio of read depth at peaks (C.C sparse/C.C abundant) respectively. This relationship was modelled across all contiguous genomic regions identified by the HMM, including both biased and unbiased regions. ‘%Motif Increase’ refers to the percentage increase in peaks containing at least one motif match in experiment biased (closed chromatin) regions relative to unbiased regions. ‘%ChIP-Signal Score Increase’ refers to the percentage increase in signal score (see Methods and Materials).

| **Cell Line** | **Amplifications Median %** | **Deletions Median %** |
| --- | --- | --- |
| K562 (Experiment Sets) | 0.139791036 | 0.134329144 |
| HepG2 (Experiment Sets) | 0.204155718 | 0.035512497 |
| GM12878 (Experiment Sets) | 0.000829876 | 0.001568422 |
| K562 (Replicate Sets) | 0.128944381 | 0.108046559 |
| HepG2 (Replicate Sets) | 0.223114957 | 0.024917442 |
| GM12878 (Replicate Sets) | 0.000963882 | 0.001364445 |

**Supplementary Table 2:**

The median % of peaks in the most biased state that reside within an identified structural variant for experiment and replicate sets in each cell line.

| **Comparison** | **Slope** | **R^2^** | **P-value** | **Adjusted P-value** |
| --- | --- | --- | --- | --- |
| (Replicate Sets) Motif Enrichment vs Deletions | 0.081 | 0.067 | 0.860 | 1 |
| (Replicate Sets) Bias Severity vs Deletions | -0.023 | 0.014 | 0.092 | 0.827 |
| (Experiment Sets) Motif Enrichment vs Deletions | 0.080 | 0.028 | 0.217 | 1 |
| (Experiment Sets) Bias Severity vs Deletions | 0.001 | 7.183e-05 | 0.911 | 1 |
| (Replicate Sets) Motif Enrichment vs Amplifications | 0.162 | 0.1245 | 0.012 | 0.127 |
| (Replicate Sets) Bias Severity vs Amplifications | -0.061 | 0.044 | 0.005 | 0.055 |
| (Experiment Sets) Motif Enrichment vs Amplifications | 0.021 | 0.001 | 0.725 | 1 |
| (Experiment Sets) Bias Severity vs Amplifications | -0.019 | 0.007 | 0.458 | 1 |
| (Replicate Sets) Motif Enrichment vs GC Content | -0.985 | 0.018 | 0.320 | 1 |
| (Replicate Sets) Bias Severity vs GC Content | 0.283 | 0.0033 | 0.543 | 1 |
| (Experiment Sets) Motif Enrichment vs GC Content | -2.1424 | 0.070 | 0.0618 | 0.618 |
| (Experiment Sets) Bias Severity vs GC Content | 0.035 | 0.000 | 0.8773 | 1 |

**Supplementary Table 3: Table of results for testing the association of structural variant overlap & GC content with motif enrichment and bias severity:**

Structural variant overlap was quantified as the proportion of peaks within a given state that overlap with a region annotated as containing a structural variant (amplification or deletion). Associations were measured between all pairs of states within the same experiment or replicate set. P values were calculated via a permutation test; The values of each variable were randomly reassigned across states in the same experiment or replicate set, and the slope coefficient for each comparison was recorded for each of n=10,000 permutations to construct an empirical null distribution. P values were corrected for multiple comparisons using the Holm-Bonferroni method^34^.

| **Comparison** | **Slope** | **P-value** | **Adjusted P-value** |
| --- | --- | --- | --- |
| (Experiment Sets) Uniformity vs Bias Occurrence | 0.162 | 0.774 | 1 |
| (Experiment Sets) Cross Correlation vs Bias Occurrence | -0.261 | 0.033 | 0.401 |
| (Experiment Sets) Enrichment vs Bias Occurrence | 1.21 | 0.047 | 0.518 |
| (Experiment Sets) Read Depth vs Bias Occurrence | -2.37E-07 | 0.260 | 1 |
| (Experiment Sets) Uniformity vs Bias Severity | 0.071 | 0.003 | 0.051 |
| (Experiment Sets) Cross Correlation vs Bias Severity | 0.019 | 0.104 | 0.938 |
| (Experiment Sets) Enrichment vs Bias Severity | 0.010 | 0.011 | 0.158 |
| (Experiment Sets) Read Depth vs Bias Severity | -1.80E-09 | 0.892 | 1 |
| (Replicate Sets) Uniformity vs Bias Occurrence | -0.006 | 0.972 | 1 |
| (Replicate Sets) Cross Correlation vs Bias Occurrence | -0.086 | 0.060 | 0.603 |
| (Replicate Sets) Enrichment vs Bias Occurrence | 0.096 | 0.004 | 0.064 |
| (Replicate Sets) Read Depth vs Bias Occurrence | -2.03E-07 | 0.012 | 0.158 |
| (Replicate Sets) Uniformity vs Bias Severity | 0.015 | 0.316 | 1 |
| (Replicate Sets) Cross Correlation vs Bias Severity | 0.001 | 0.830 | 1 |
| (Replicate Sets) Enrichment vs Bias Severity | 0.003 | 0.315 | 1 |
| (Replicate Sets) Read Depth vs Bias Severity | -3.73E-09 | 0.653 | 1 |

**Supplementary Table 4: Table of results for testing the association between ChIP-seq quality metrics and the occurrence and severity of regional bias.**

For each quality metric, logistic regression was used to model the association between the minimum quality score across replicates (for replicate sets) or experiments (for experiment sets) and the probability of regional bias occurring within each replicate or experiment set. Linear regression was used to model the relationship between the between the minimum quality score across replicates (for replicate sets) or experiments (for experiment sets) and the maximum bias severity observed across states normalised to the average bias severity of all states within a given experiment set. The R package ChIPQC^27^  was used to calculate quality scores, and P values were corrected for multiple comparisons using the Holm-Bonferroni method^34^.
